# scRGP: Prediction of Single-cell Genetic Perturbation Transcriptional Responses based on Rank in Multiple Scenarios

**DOI:** 10.1101/2025.09.30.679477

**Authors:** Yufeng Liu, Hao Zhang, Mingmin Xu, Duowei Wang, Weiwei Hu, Liangyun Zhang, Yong Yang, Cong Pian, Yuanyuan Chen

**Affiliations:** College of Agriculture, Nanjing Agricultural University, Nanjing, Jiangsu 210095, China; College of Sciences, Nanjing Agricultural University, Nanjing, Jiangsu 210095, China; School of Basic Medicine Sciences, Gannan Medical University, Ganzhou, Jiangxi 341000, China; School of Basic Medicine and Clinical Pharmacy, China Pharmaceutical University, Nanjing, Jiangsu 211198, China; Institute of Translational Medicine, China Pharmaceutical University, Nanjing 210009, China

**Author notes:** Corresponding authors: Email adress. These authors contributed equally to this work (Yufeng Liu, Hao Zhang, Mingmin Xu).

## Abstract

Single-cell perturbation sequencing technologies (e.g., Perturb-seq, CROP-seq), which integrate CRISPR-based gene editing with single-cell transcriptome profiling, have revolutionized the analysis of transcriptomic changes induced by genetic perturbations at single-cell resolution. These technologies serve as a powerful tool for identifying key genes that inhibit tumor growth or reverse cancer cell phenotypes. However, they face two major challenges: data explosion with high experimental costs, and data complexity characterized by high dimensionality, noise, sparsity, and heterogeneity. To address these challenges, we developed the single-cell Rank-based Genetic Perturbation predictor (scRGP), the first deep learning framework leveraging gene expression rank-order information for this task. scRGP demonstrates superior performance in terms of robustness, cross-cell-line perturbation prediction, and high-throughput screening. Specifically, scRGP achieves an approximately 10-16 percentage points improvement in Pearson correlation coefficient (PCC) over state-of-the-art methods (e.g., GEARS and scFoundation) for single- and double-gene perturbation predictions, while also extending prediction capability to triple-gene perturbations. Furthermore, it outperforms these methods by approximately 5-9 percentage points in cross-cell-line predictions. These advancements promise to shift the paradigm of single-cell perturbation studies from experiment-driven to computation-driven approaches, providing new support for functional genomics and precision medicine.

## Introduction

Breakthroughs in single-cell transcriptome sequencing technologies, particularly the development of single-cell perturbation sequencing methods such as Perturb-seq^[1]^ and CROP-seq^[2]^, have provided powerful tools for systematically deciphering gene function and regulatory networks^[1, 3, 4]^. With increasing experimental throughput and accuracy, large-scale single-cell perturbation transcriptomic datasets have been generated^[2, 5-9]^, laying a solid data foundation for computational prediction of gene perturbation effects and global gene interaction mapping. However, the vast scale of the human genome and the diversity of cell types make it infeasible to exhaustively explore all genes or even combinatorial perturbations experimentally^[10, 11]^. Therefore, developing computational methods capable of accurately and generalizably predicting transcriptional responses to arbitrary genetic perturbations has become a central challenge in revealing gene function and regulatory networks and advancing precision medicine^[12-14]^.

The key scientific problem lies in constructing a predictive model unconstrained by the lack of prior knowledge, while still achieving strong generalization across unseen genes and cell types. Although existing methods have made valuable contributions, they exhibit notable limitations: models such as GEARS heavily rely on structured biological knowledge (e.g., GO annotations) and fail to predict perturbations for unannotated genes^[10]^; VAE-based approaches (e.g., scGEN^[14]^, CPA^[13]^) are highly sensitive to training data distribution and exhibit limited generalization; optimal transport methods (e.g., CellOT^[15]^) underperform in cross-cell-type prediction tasks; and large pre-trained models (e.g., Geneformer^[16]^, scGPT^[17]^) suffer from high computational demands and requirements for customized input representations^[18-20]^. Additionally, the inherent high noise, sparsity, and batch effects in scRNA-seq data further complicate the development of robust models^[21, 22]^.

In response to these challenges, we propose scRGP (single-cell Rank-based Genetic Perturbation predictor), a deep learning framework leveraging gene expression ranking structures. Our approach eliminates reliance on external prior knowledge and instead capitalizes on the robust biological signals inherent in gene expression rankings — a strategy validated by pioneering works such as Geneformer and GeneCompass, which effectively mitigate noise and batch effects^[16, 23-25]^. By constructing a multilayer neural network that directly learns latent gene-gene and cell-state relationships from ranked data, scRGP enables end-to-end perturbation response prediction without predefined gene embeddings or functional annotations.

Experimental results demonstrate that scRGP achieves significant breakthroughs across multiple key scenarios: it not only excels in predicting transcriptional responses to unseen genetic perturbations but also generalizes efficiently to entirely novel cell types. Its performance consistently surpasses that of leading baseline and foundation models, underscoring the advantage of rank-based representations in genetic perturbation prediction. Our study provides an efficient, lightweight, and versatile tool for predicting perturbation effects while also offering a new paradigm for inferring global gene functional interactions from limited data.

## Results

### Method overview

Figure 1 presents scRGP, a deep learning-based predictor of single-cell perturbation transcriptional responses, which forecasts post-perturbation transcriptional profile ranks for individual or combinatorial perturbations (see Methods). The framework assigns each gene a unique embedding as its core feature, which is integrated with perturbed gene embeddings during training to model gene-gene interaction effects. These integrated embeddings undergo dimensionality reduction to generate vectors matching the dimensionality of cellular transcriptional profiles, serving as the primary input for model training. Concurrently, pre-perturbation transcriptional profiles provide secondary input by converting gene expression data into rank-based vectors, thereby amplifying the contribution of highly expressed genes during model learning.

**Figure 1:**
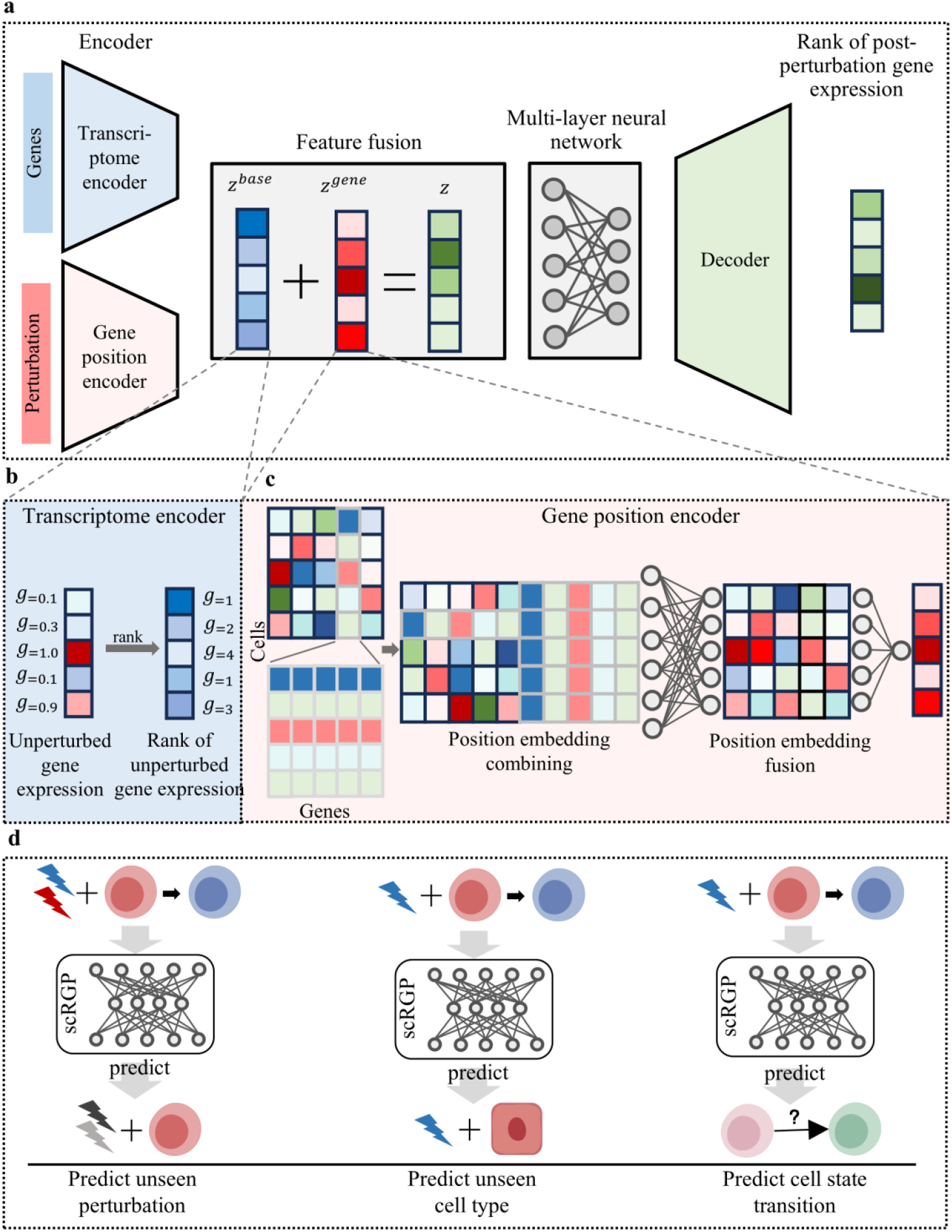
Overview of the methodological approach. **a**. scRGP uses rank-based gene expression to predict post-perturbation gene expression ranks. **b**. Transcriptome encoding module. **c**. Gene location encoding module. **d**. Downstream tasks implemented by scRGP: the left panel predicts unknown perturbations (including single-gene and multi-gene perturbations), the middle panel predicts perturbations in unknown cell types, and the right panel predicts cell state changes.

scRGP enables multiple predictive tasks including single-gene perturbation transcription prediction, multi-gene perturbation transcription prediction (encompassing dual-gene and triple-gene combinations), and cross-cell-type prediction. By consolidating diverse functionalities within a streamlined architecture, the model demonstrates robust scalability and versatility.

### scRGP predicts transcriptional responses to single-gene perturbations

In single-gene perturbation transcriptional prediction, scRGP partitions datasets by perturbed genes, ensuring test datasets remain previously unseen during model training (Figure 2a). Five genetic perturbation datasets from independent screening studies were utilized: 89 perturbations from Adamson et al.^[26]^, 23 from Dixit et al.^[1]^, 231 from Norman et al. (including 100 single-gene and 131 double-gene perturbations)^[27]^, and 1,756 (RPE1 cells) plus 1,751 (K562 cells) perturbations from Replogle et al.^[4]^ (Supplementary Table 1; Methods). These datasets are hereafter referred to as Adamson, Dixit, Norman, Replogle_RPE1, and Replogle_K562. scRGP was trained on each dataset, alongside two comparative models: GEARS^[10]^, an existing deep learning approach, and a baseline model assuming no transcriptomic effect from perturbations, plus scFoundation^[28]^ (Methods).

**Figure 2:**
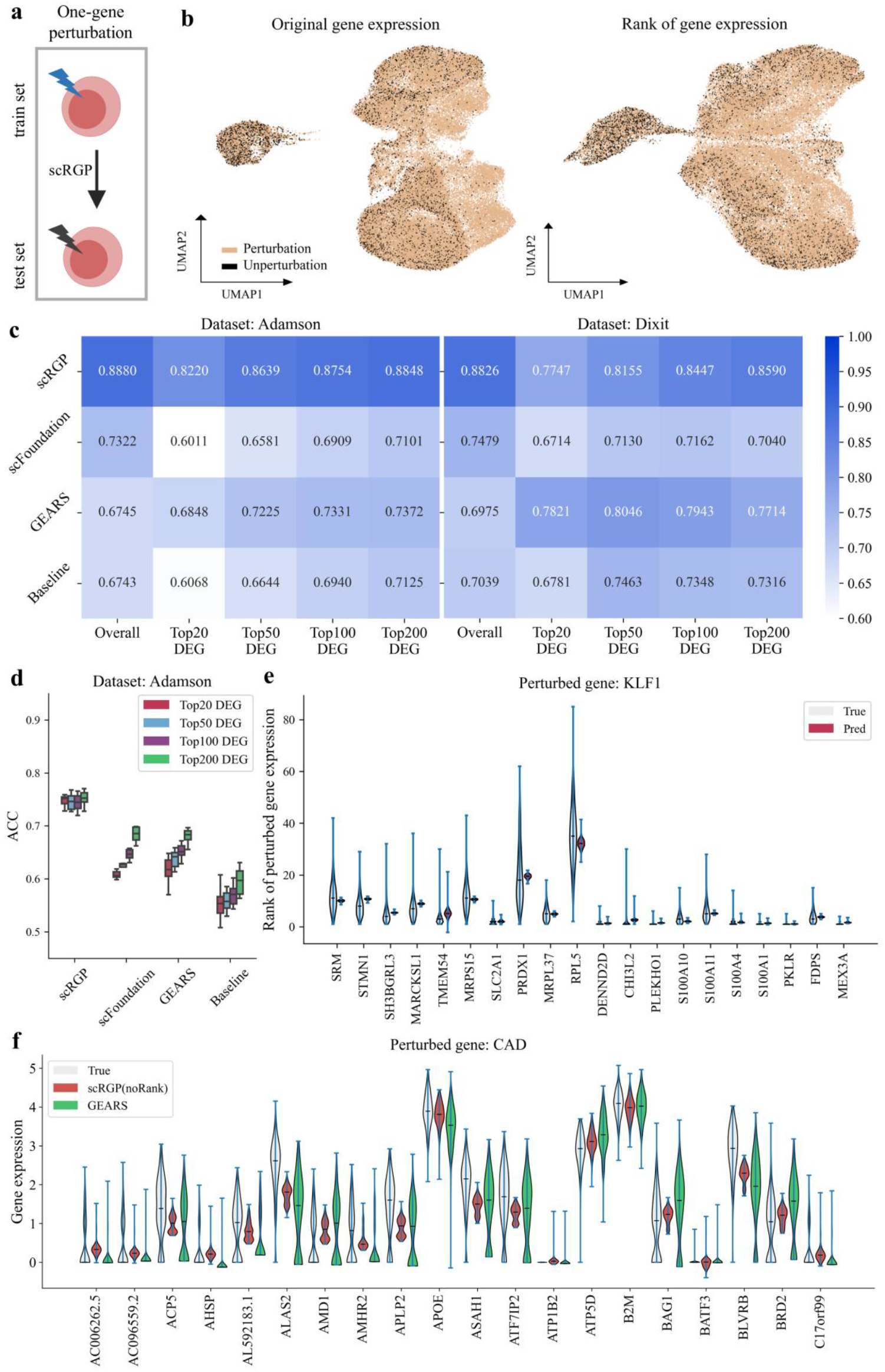
The ability of scRGP to predict the ranks of post-perturbation gene expression profiles under single-gene perturbation. **a**. Training and testing data splitting scheme for single-gene perturbation; **b**. UMAP projections of Adamson dataset showing: original unperturbed vs. post-perturbation gene expression, and pre-vs. post-perturbation rank-based gene expression; **c**. Pearson correlation coefficients of the four models in Adamson and Dixit datasets; **d**. Accuracy of DEG expression direction changes post-perturbation in Adamson across the four models; **e**. Violin plots comparing actual vs. scRGP-predicted gene expression for the top 20 DEGs following KLF1 perturbation in the Norman dataset; **f**. Violin plot showing true gene expression, scRGP-predicted expression (without Rank module), and GEARS-predicted expression for the top 20 DEGs following CAD perturbation in the Adamson dataset.

Performance evaluation employed Pearson correlation coefficients (PCC) to compare actual versus predicted post-perturbation expression and accuracy (ACC) to assess the direction of gene expression changes. Final metrics represent averages across all cell pairs (see Methods). UMAP visualization confirmed minimal differences between original and rank-based gene expression distributions (Figure 2b). Overall, scRGP exhibited superior predictive performance, achieving the highest PCC for global gene expression predictions—outperforming the baseline model by approximately 18-28%. This indicates effective capture of underlying biological patterns through rank-based expression. In contrast, GEARS performed comparably to the baseline, while scFoundation—though integrating transcriptomic and perturbation information into holistic embeddings—still showed 10-16% lower PCC than scRGP for perturbation-induced expression predictions (Figure 2c; Supplementary Figure 1a).

Given the sparsity of single-cell transcriptomes—where only a subset of genes correlates strongly with perturbed genes—we analyzed PCC and ACC for the top 20, 50, 100, and 200 differentially expressed genes (DEGs). scRGP consistently outperformed other models in DEG PCC except for the top 20 DEGs in Dixit (≤1% difference from GEARS). Notably, scRGP achieved >20% PCC improvements over the baseline across all DEG categories in Adamson. In Dixit, while scRGP showed slightly lower PCC than GEARS for top 20 DEGs, it outperformed by 1%, 5%, and 8% for top 50, 100, and 200 DEGs respectively. scFoundation outperformed GEARS in Replogle_RPE1/Replogle_K562 but underperformed in Adamson/Dixit/Norman, potentially due to its emphasis on global latent information over specific gene effects (Figure 2c; Supplementary Figure 1a). For ACC, scRGP showed moderate performance (70-80% across datasets), with strongest results in Adamson and parity with top-performing GEARS in Norman, but lower performance in other datasets (Figure 2d; Supplementary Figure 1b). This may stem from the expanded value range introduced by rank-based expression, which could diminish sensitivity for predicting expression directionality.

To gain clearer insight into the model’s predictions versus actual post-perturbation gene expression variations, we randomly selected a perturbed gene (KLF1) from the Norman dataset and analyzed the expression of the top 20 DEGs (Figure 2e). As observed, rank-based gene expression values following actual perturbation exhibit substantial variability across cells, whereas scRGP predictions display stability. The medians of actual and predicted values are closely aligned, confirming the predictive accuracy of our model.

In addition, we briefly evaluated the predictive performance of the model with the rank module removed (Figure 2f; Supplementary Note 1). Specifically, in the Adamson dataset, we selected the top 20 DEGs observed following perturbation of the CAD gene and compared their expression profiles with predictions from GEARS. scRGP’s predictions showed closer alignment with actual gene expression levels (Figure 2f).

### scRGP predicts transcriptional responses to multiple-gene perturbations

scRGP is designed to predict transcriptional outcomes of perturbation groups composed of multiple gene combinations. For two-gene perturbation combinations, we evaluated model performance using the Norman dataset. Three data splitting strategies were defined in this scenario (Figure 3a): (i) neither gene in the combination was included in training (zero seen of two), (ii) one of the two genes had been trained (one seen of two), and (iii) both genes in the combination were part of the training set (two seen of two).

**Figure 3:**
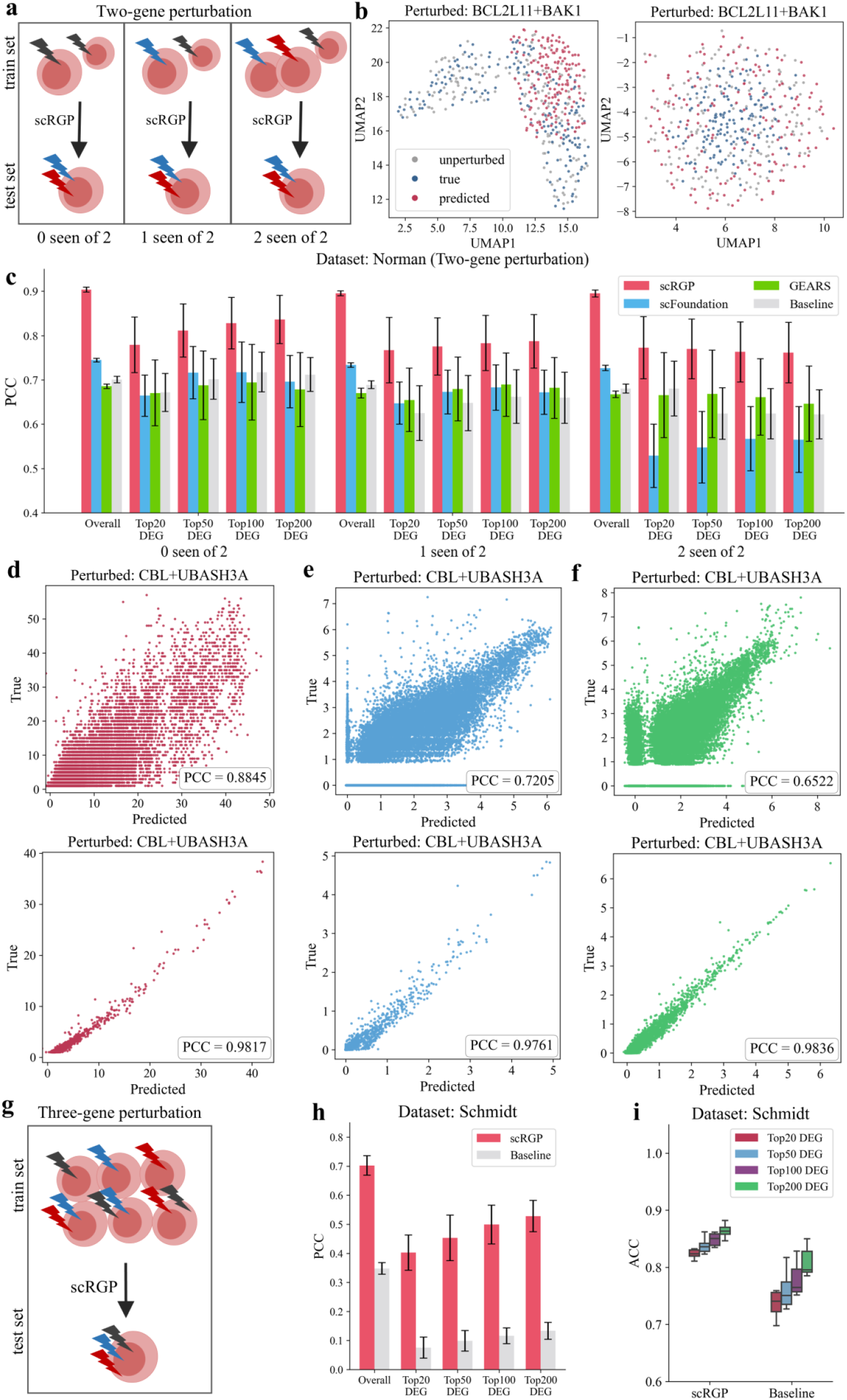
The ability of scRGP to predict the ranks of post-perturbation gene expression profiles under two-gene perturbations. **a**. Training and testing data splits for two-gene perturbation scenarios; **b**. UMAP projections showing predicted expression profiles of the BCL2L11+BAK1 perturbation combination under the ‘1 seen of 2’ condition (scRGP predictions on left, GEARS on right); **c**. Pearson correlation coefficients comparing predicted vs. actual post-perturbation gene expression for scRGP, scFoundation, GEARS, and Baseline models; **d-f**. Scatter plots of predicted vs. actual post-perturbation gene expression for scRGP (d), scFoundation (e), and GEARS (f). Top row: one-to-one cell matching; Bottom row: mean expression values across all cells per gene; g. Training and testing data split strategy for three-gene perturbation prediction; **h**. Pearson correlation coefficient of scRGP on the Schmit dataset; **i**. Accuracy comparison for predicting top 20, 50, 100, and 200 DEGs in Schmit’s perturbation datasets.

scRGP demonstrated superior predictive performance across all three scenarios; it outperformed the next best model by over 16% for all genes and 10% for DEGs, respectively (Figure 3c). Specifically, for all-gene expression prediction, scRGP’s performance remained largely consistent across the three data splits, whereas scFoundation showed slightly lower performance. GEARS exhibited predictive capability comparable to the baseline model. However, for DEGs, scFoundation and GEARS showed nearly opposite performance trends: scRGP’s DEG prediction performance gradually deteriorated across the zero-seen, one-seen, and two-seen categories. This may result from cumulative negative effects introduced by progressively incorporating single-gene perturbations from combination perturbations during training, which increases prediction complexity.

We further analyzed results at the individual perturbation level. For BCL2L11/BAK1 combined perturbation prediction, UMAP projections showed partial overlap between scRGP-predicted results and actual post-perturbation cellular expression profiles, whereas GEARS-predicted cells clustered around unperturbed cells (Figure 3b). Scatter plots of predicted expression levels for CBL/UBASH3A combined perturbation revealed striking performance differences among the three models under one-to-one cell matching (top row, Figure 3d, 3e, and 3f). However, when calculating mean expression values across all cells for each gene, all models exhibited strong performance with similar results (bottom row, Figure 3d, 3e, and 3f). This is because averaging stabilizes expression levels, thereby enhancing evaluation metrics like Pearson correlation coefficients. Nevertheless, average values no longer represent single-cell transcriptional profiles and fail to capture the accuracy of single-cell perturbation response predictions. For this reason, our model evaluation method involved computing metrics for each cell pair and averaging the results.

In addition, we conducted preliminary exploration of three-gene combination perturbation predictions. Given the large number of genes, three-gene perturbation combinations are highly diverse, but available data remain limited. We therefore employed a single data partitioning method without distinguishing whether test set perturbation genes were present in the training set (Figure 3g). Evaluation metrics showed improvements of 30-40% in Pearson correlation coefficient and 5-8% in DEG expression direction prediction accuracy (Figure 3h, 3i).

### scRGP predicts transcriptional responses to perturbations in unseen cell types

Here, we established a benchmark dataset using Replogle_K562 and Replogle_RPE1 datasets, identifying 1,448 overlapping perturbation genes and 3,757 shared genes. For this cross-cell-type prediction task, we first trained the model on Replogle_K562 data to predict post-perturbation expression ranks in Replogle_RPE1, followed by the reverse procedure (Figure 4a).

**Figure 4:**
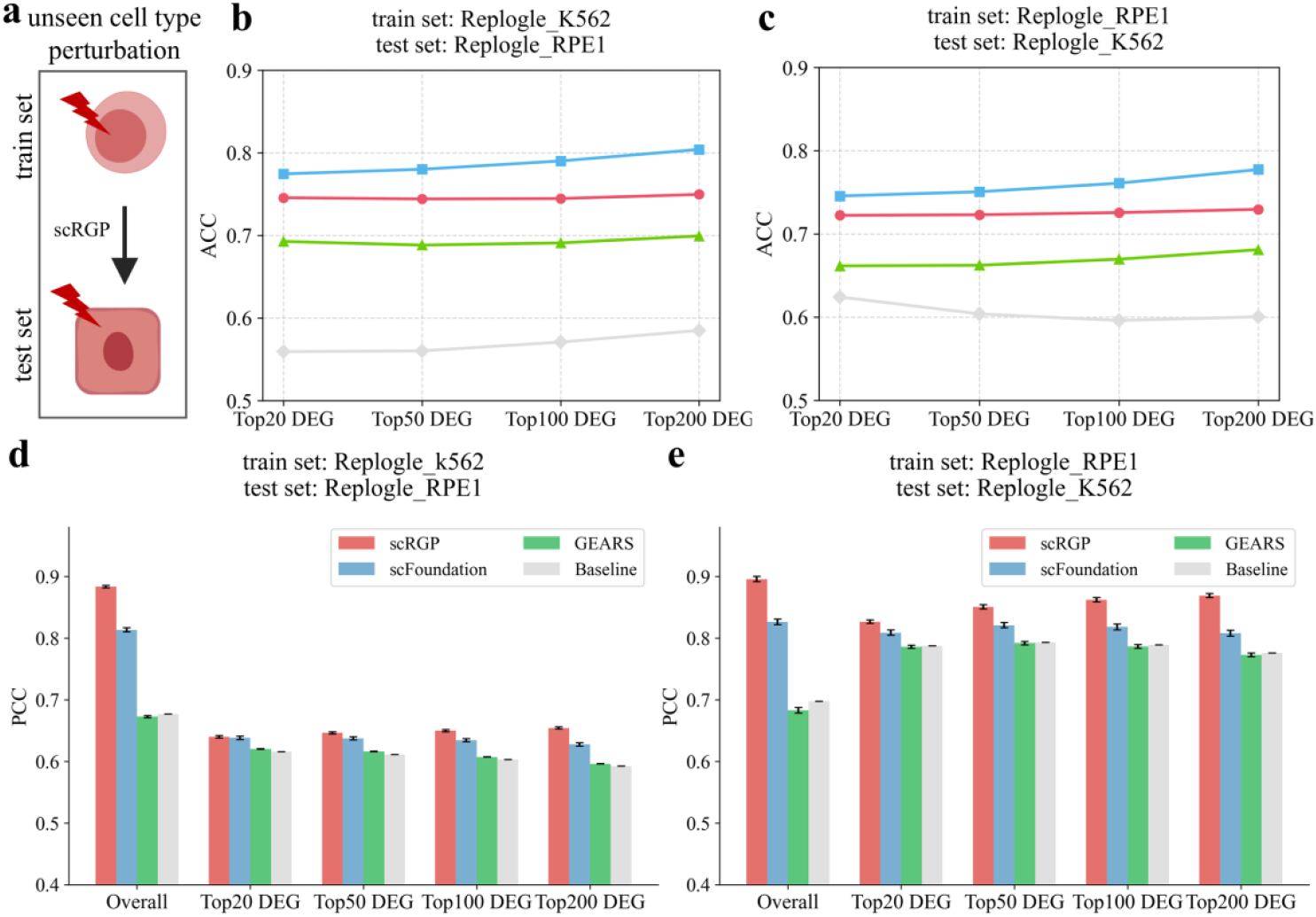
The ability of scRGP to predict cross-cell-type transcriptional responses to unseen cell lines. **a**. Training/testing set splits for unseen cell type prediction; **b**. Accuracy of Replogle_K562-trained models in predicting DEG expression direction changes in Replogle_RPE1; **c**. Accuracy of Replogle_RPE1-trained models in predicting DEG expression direction changes in Replogle_K562; **d**. Pearson correlation coefficients for Replogle_K562-trained models predicting Replogle_RPE1; **e**. Pearson correlation coefficients for Replogle_RPE1-trained models predicting Replogle_K562.

scRGP demonstrated superior predictive performance under both test conditions. When trained on Replogle_K562 to predict post-perturbation gene expression in Replogle_RPE1, scRGP achieved a Pearson correlation coefficient of 0.89 across all genes. However, its DEG prediction performance was modest, showing only a 3% improvement over the baseline for top 20 DEGs, with gradual enhancement as the number of DEGs increased (Figure 4d). This discrepancy may result from scRGP’s higher accuracy in predicting low-expression genes, leading to a significantly higher Pearson correlation coefficient for all genes compared to DEGs. Conversely, when trained on Replogle_RPE1 to predict perturbations in Replogle_K562, scRGP exhibited robust performance—outperforming the baseline by ∼20% across all genes and 4-9% for top 20-200 DEGs. As expected, GEARS showed strong cell line specificity due to its reliance on prior knowledge, resulting in minimal predictive capability on unseen cell types (performance comparable to baseline). scFoundation showed moderate improvement over GEARS in this scenario (Figure 4e), and additionally outperformed scRGP by 2.5-5% in predicting gene expression direction changes (Figure 4b-c).

### scRGP predicts cell state transitions

In biological processes such as development, changes in cell function or state are driven by single genes or gene groups. Here, we utilized the single-cell dataset Ainciburu^[29]^, which demonstrates that SPI1 promotes lineage changes in granulocyte-monocyte progenitors (GMPs), while GATA1 induces transformation of megakaryocyte-erythroid progenitors (MEPs); specifically, SPI1 knockout converts GMPs to lymphoid-myeloid progenitor cells (LMPPs), and GATA1 knockout converts MEPs to LMPPs^[19]^.

We generated UMAP projections to visualize cellular distributions (Figure 5a), pre-perturbation cell state transition directions (Figure 5b), and predicted post-perturbation transition directions (Figures 5c, 5d). Cell state transition directions were visualized using streamlines. When predicting cellular transition trends following SPI1 knockout, both scRGP and GEARS partially captured directional changes. However, for GATA1 knockout-induced changes, GEARS failed to capture lineage-specific transitions, whereas scRGP retained partial accuracy. Overall, predicting cell state transitions remains a highly challenging task.

**Figure 5:**
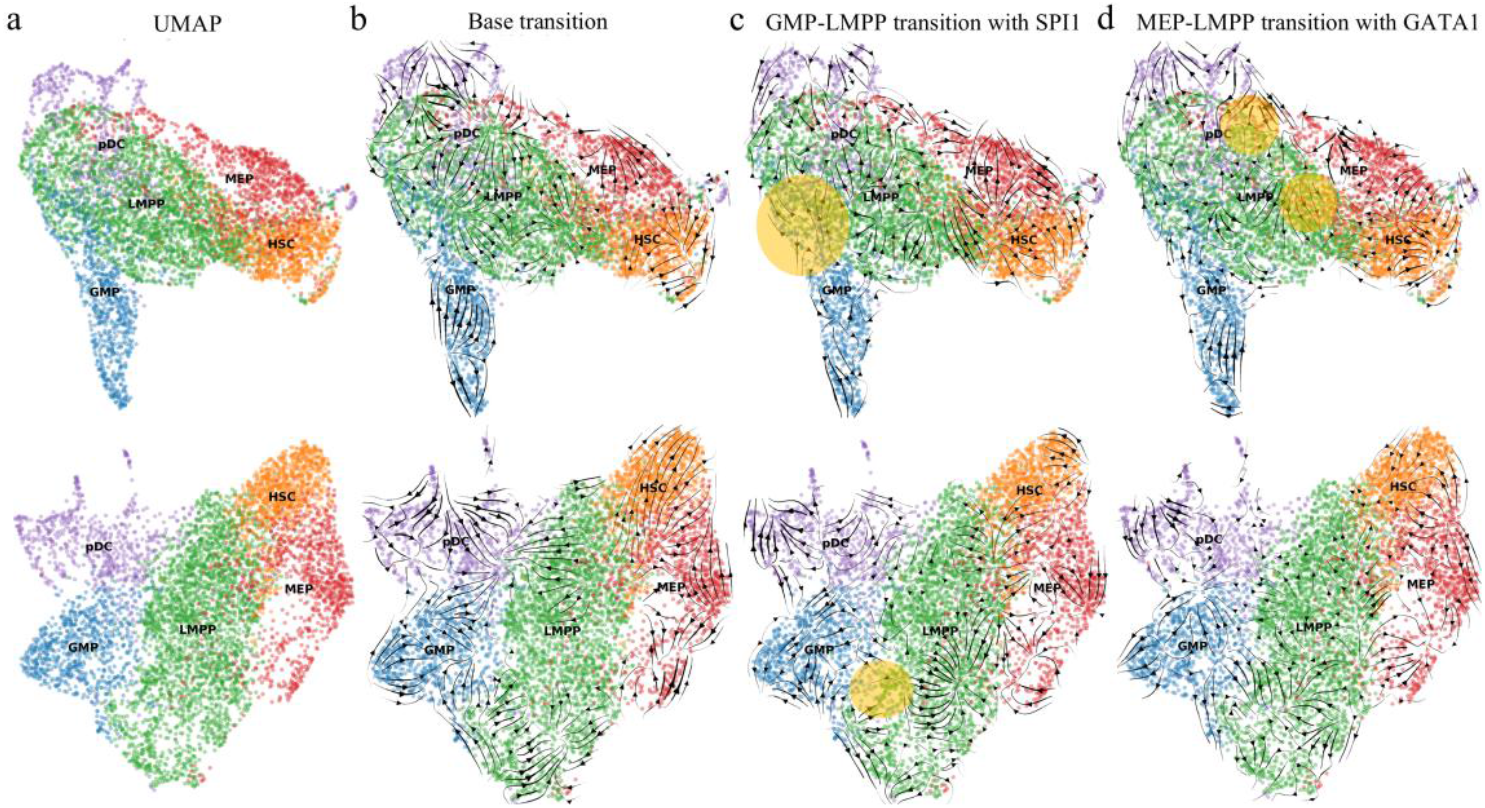
Test of scRGP predicting cell state transitions. **a**. UMAP projections of five cell types; **b**. Pre-perturbation cell trajectory; **c**. GMP-to-LMPP transition following SPI1 knockdown; **d**. MEP-to-LMPP transition following GATA1 knockdown. **a-d**. Top row: scRGP-predicted post-perturbation UMAP projections of gene expression ranks with streamline trajectories; Bottom row: GEARS-predicted post-perturbation UMAP projections of gene expression with streamline trajectories.

## Discussion

In view of the two key problems in predicting transcriptional responses to genetic perturbations—namely, the inherent high noise, sparsity, and batch effects of scRNA-seq data, and the poor cross-scenario predictive capability of existing models—we propose a rank-based genetic perturbation prediction method called scRGP. Our study presents the innovations and advantages of scRGP, along with several key highlights that advance the field of single-cell perturbation prediction:

### 1. Robust performance across perturbation types

scRGP not only demonstrates superior predictive accuracy for single-gene perturbations (achieving 10-16% higher PCC than state-of-the-art methods) but also extends this capability to multi-gene combinations — including the first model to successfully predict three-gene perturbation effects (30-40% PCC improvement). This breakthrough addresses a critical gap in current computational tools, which are limited to single- or double-gene predictions.

### 2. Unprecedented cross-cell-type generalization

The model achieves remarkable transfer learning performance (PCC=0.89) when predicting responses in completely unseen cell types (K562→RPE1 and vice versa), outperforming existing methods by 5-9 percentage points. This suggests that rank-based representations effectively capture conserved biological patterns across cellular contexts.

### 3. Computational efficiency without sacrificing accuracy

Despite using a lightweight MLP architecture (1% of the parameters of scGPT), scRGP maintains competitive performance while enabling rapid screening. This makes it particularly suitable for large-scale virtual screening scenarios where computational resources are limited.

### 4. Novel biological insights

scRGP can reveal potential novel regulatory relationships through its perturbation predictions. Its capability to predict cell state transitions (e.g., GMP→LMPP transition upon SPI1 knockout) further demonstrates its biological relevance.

These advances carry significant implications for both basic research and translational applications: (i) For fundamental science, scRGP provides a new paradigm for exploring gene regulatory networks at scale, particularly for studying complex multi-gene interactions that are experimentally challenging to probe^[30, 31]^. The success of rank-based representations suggests gene expression hierarchies may encode more fundamental biological information than absolute expression levels; (ii) For discovery of target combinations, the model’s ability to prioritize effective perturbation combinations (including triple-gene targets) could accelerate identification of therapeutic candidates, especially for complex diseases requiring combinatorial interventions. Its cross-cell-type predictions may help bridge gaps between model systems and human biology; (iii) For experimental design, by reducing reliance on prior knowledge, scRGP enables exploration of understudied genes and cell types, potentially uncovering novel biology missed by annotation-dependent approaches.

Moving forward, we propose several promising research avenues: (i) incorporating temporal dynamics to model perturbation response trajectories; (ii) extending to multi-omic perturbations, including epigenetic modifications; and (iii) developing interpretability tools to extract mechanistic insights from predictions. While current limitations around biological interpretability and cell cycle effects warrant further investigation^[32-34]^, scRGP establishes a versatile foundation for computational perturbation biology that balances performance, generality, and scalability.

## Methods

### Overview of scRGP

scRGP requires a perturbation dataset 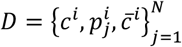 consisting of multiple cells, where *c*^*i*^ ∈ ℂ^*K*^ represents the expression vector of *K* genes in cell *i* before perturbation, 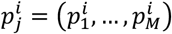 represents the *M* perturbation sets in cell *i*, and 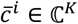 represents the expression vector of *K* genes in cell *i* after perturbation, the perturbed cell *i* is randomly matched with the unperturbed cell *i*. Each gene *K* in the ℂ^*K*^ dataset has a unique embedding as its identifier. scRGP ultimately learns a function *f* that maps perturbation sets and pre-perturbation cell transcriptomes to post-perturbation results, i.e., gene expression 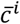.

### Gene position coding and gene expression coding

To set up gene identifiers, scRGP randomly embeds each gene 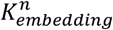, representing the embedding on *n* = (1,2, … *n*) genes, perturbs gene 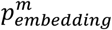 by taking out its corresponding position m perturbed gene embedding, and then combines all genes and perturbed gene embeddings into 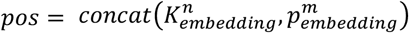. The training process is *MLP*(*pos*), followed by fusion layer *f*_*pos*_ = *MLP*{*MLP*(*pos*)}.

We process gene expression values into ranked representations *g*^*R*^ = *Rank*(*g*) to increase the importance of highly expressed genes in model training, namely *f*_*gene*_ = *g*^*R*^.

### Train and test

The specific process is as follows: given a perturbed dataset, there exists a perturbation set *p* = (*p*_1_, …, *p*_*M*_), where each perturbation has multiple cells 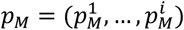. scRGP first assigns a unique position encoding to each gene within a cell, then concatenates all gene position encodings with the perturbation gene encodings and maps them through a neural network to a K-dimensional gene position embedding

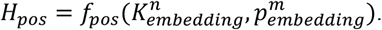

Next, it maps the pre-perturbation gene expression to a rank-based expression

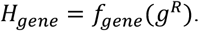

Finally, combine the gene position embedded in *f*_*pos*_ and gene expression *f*_*gene*_ into

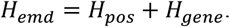

And input into a multi-layer neural network for learning, then mapped through a decoder

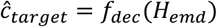

to the rank of gene expression after perturbation.

### Loss function

In their work on GEARS, the authors proposed a new composite loss function and verified that its effectiveness greatly exceeds that of the MSE loss function. Therefore, we borrowed the loss function defined by GEARS, namely

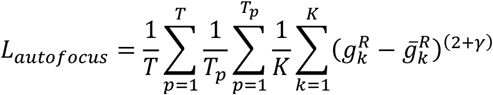

Given a perturbation set *T*, where each perturbation p has *T*_*p*_ cells with *K* genes, it has predicted post-perturbation gene expression 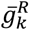 and true gene expression 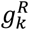 . Here, we set *γ* to 2.

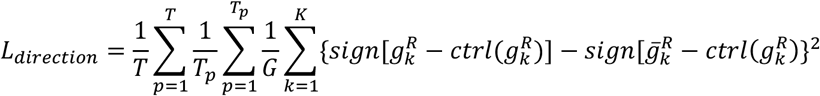

The predicted loss function is *L* = *L*_*autofocus*_ + *λ*L_*direction*_, where *λ* is set to 0.001.

### Data process

Perturbation data processing: Cells with fewer than 30 perturbations were excluded; however, due to limited three-gene perturbation data, cells with two or more perturbations were retained. Low-quality cells expressing fewer than 200 genes were removed. The top 5,000 highly variable genes were retained, along with perturbation genes not included in this set. Normalization was performed using Seurat v3 via Scanpy^[35]^. For model training, datasets were split into training, validation, and test sets at an 8:1:1 ratio.

Cross-cell line analysis: Intersection genes and overlapping perturbed genes were selected from processed Replogle_RPE1 and Replogle_K562 datasets. Models were trained sequentially on one dataset and then used to predict perturbation responses in the other.

Cell state transition data processing: Following Chen et al.^[19]^, highly variable genes were selected using Scanpy with parameters: min_mean=0.0125, max_mean=3, and min_disp=0.4.

Detailed dataset information is provided in Supplementary Table 1.

### Baseline models

We employed the following baseline models for performance comparison:

#### Baseline

To verify genuine improvements in model performance, we directly computed the correlation between pre-perturbation and actual post-perturbation gene expression, which served as the baseline.

#### GEARS^[10]^

A deep learning model enhanced by prior biological knowledge. The original evaluation method involved averaging predictions across multiple cells for the same perturbation before computing metrics. Here, we modified this approach by matching each perturbation to its corresponding cell, calculating metrics for individual predicted cells, and then averaging the results.

#### scFoundation^[28]^

A foundation model that integrates perturbation and transcriptomic data to generate comprehensive embeddings, requiring downstream models to be trained on these embeddings for prediction. In the original study, GEARS was used as the downstream model, with authors demonstrating that scFoundation significantly enhanced predictive performance. We adopted this same framework. Since the original evaluation metrics were calculated identically to GEARS, we similarly switched to a cell-matched calculation method.

### Evaluation metrics

Model performance was evaluated using two metrics: Pearson correlation coefficient (PCC) and directional accuracy of gene expression changes (ACC).

PCC is calculated using the following formula:

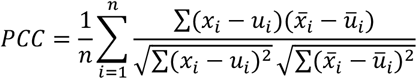

Where *x*_*i*_ and 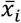 represent the actual and predicted gene expression of the cell *i* after perturbation, respectively, and *u*_*i*_ and *ū*_*i*_ represent the mean values of *x*_*i*_ and 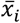, respectively. When calculating *PCC*_*DEGs*_, the gene expression of the top 20, 50, 100, and 200 DEGs is calculated, respectively.

ACC is calculated using the following equation:

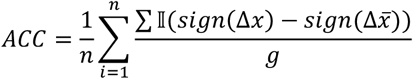

Where Δ*x* represents the value of gene expression after the actual perturbation minus gene expression before the perturbation, 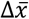 represents the predicted gene expression after the perturbation minus gene expression before the perturbation, *g* represents the number of genes, and *sign*(·) is:

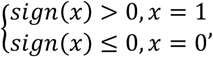

The meaning of 𝕀 (·) is

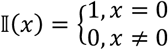

ACC values are calculated for DEGs.

### Streamline plot

Based on pre-perturbation gene expression and model-predicted post-perturbation gene expression, the direction and speed of change for each cell are simulated and calculated. Subsequently, the scVelo^[36]^ package is used to plot visualized streamlines on the UMAP.

## Data availability

All scCRISPR datasets used in scRGP are from publicly available databases: Dixit et al.^[1]^: GSE90063; Adamson et al.^[26]^: GSE90546; Norman et al.^[27]^ : GSE133344; Replogle et al.^[4]^: GSE146194; The data from Replogle et al. are available at https://doi.org/10.25452/figshare.plus.20022944. Ainciburu et al.^[29]^ : GSE180298.

## Code availability

The source code of scRGP is freely available via GitHub (https://github.com/yuf-Liu/scRGP).

## Acknowledgements

This work was supported in part by the National Natural Science Foundation of China [ZX2200521] and the Anti-tumor New Drug Rapid Translation Public Service Platform of Jiangsu Province (BM2023002).

## Contributions

Y. Chen, C. Pian and Y. Yang conceived and supervised the project. Y. Chen, H. Zhang and Y. Liu designed the experiments. H. Zhang, F. Liu and M. Xu established and validated the scRGP model. Y. Liu, D. Wang and W. Hu drafted the manuscript. Y. Chen, H. Zhang, C. Pian and L. Zhang discussed the results and revised the manuscript.

