## Supplementary information for "scRGP: Prediction of Single-cell Genetic Perturbation Transcriptional Responses based on Rank in Multiple Scenarios"

[†]The first three authors should be regarded as Joint First Authors.

**Supplementary Note 1: Ablation analysis.**

scRGP uses rank-based transcriptomic profiles to train the model. To validate the effectiveness of this module, we removed it and evaluated the model on the Adamson dataset. After removing the rank module, the model's performance significantly decreased: the Pearson correlation coefficient for all gene expression predictions decreased by approximately 10%, and the coefficient for differentially expressed genes decreased by 1–6% (Supplementary Figure 2a). However, the accuracy of predicting the direction of change in differentially expressed genes remained largely unchanged (Supplementary Figure 2b). Additionally, to validate the state changes among matched unperturbed, perturbed, and predicted perturbed cells, we generated a UMAP plot. We then randomly selected ten cell pairs and marked the matched pre-perturbation and true post-perturbation cells, as well as the pre-perturbation and predicted post-perturbation cells, with arrows (Supplementary Figure 2c). Longer arrows indicate greater differences in transcriptional profiles. We observed that after removing the rank module, the cells predicted by scRGP showed smaller differences from the true post-perturbation cells compared to the pre-perturbation cells. Furthermore, the lengths and directions of the arrows from pre-perturbation cells to true post-perturbation cells and from pre-perturbation cells to predicted post-perturbation cells (as determined by scRGP) were comparable and similar in direction.

**Supplementary Table 1: Overview of the dataset**

| Author | Dataset | Cell Type | Perturbation | Cells | Type |
|---|---|---|---|---|---|
| Adamson et al.[1] | Adamson | K562 | 89 | 64861 | One-gene |
| Dixit et al.[2] | Dixit | K562 | 23 | 64228 | One-gene |
| Norman et al.[3] | Norman | K562 | 231 | 102819 | Two-gene |
| Replogle et al.[4] | Replogle_RPE1 | RPE1 | 1756 | 207289 | One-gene |
| Replogle et al.[4] | Replogle_K562 | K562 | 1751 | 283593 | One-gene |
| Schmidt et al.[5] | Schmidt | T cell | 406 | 71841 | Three-gene |
| Ainciburu et al.[6] | Ainciburu | HSPC | | | |

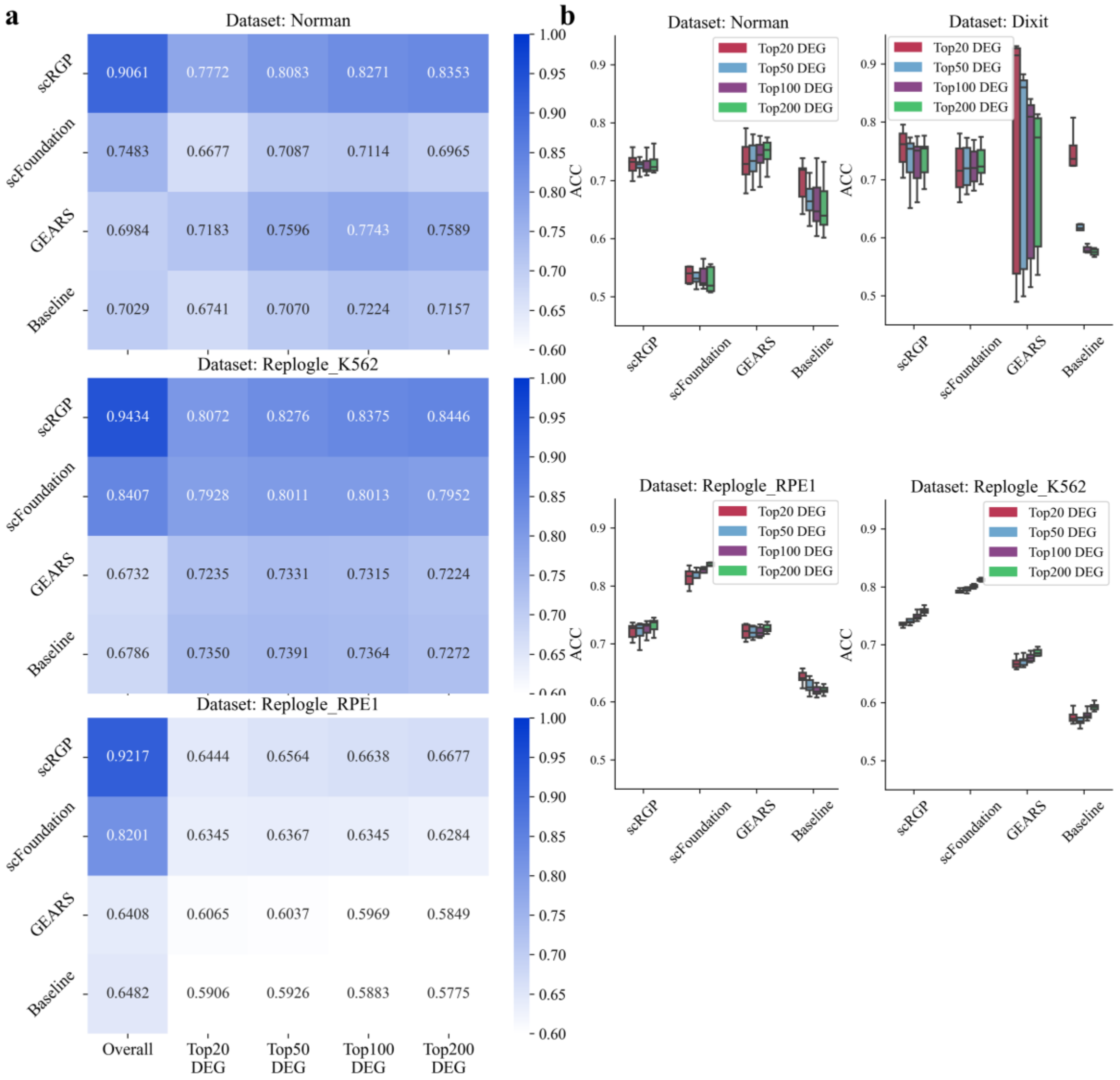


**Supplementary Figure 1: Evaluation of scRGP across multiple data sets. a.** Comparison of PCCs for all genes, top 20 DEGs, top 50 DEGs, top 100 DEGs, and top 200 DEGs in perturbation datasets from Norman, Replogle_RPE1, and Replogle_K562. **b.** Comparison of ACC for top 20 DEGs, top 50 DEGs, top 100 DEGs, and top 200 DEGs in perturbation datasets from Norman, Dixit, Replogle_RPE1, and Replogle_K562.

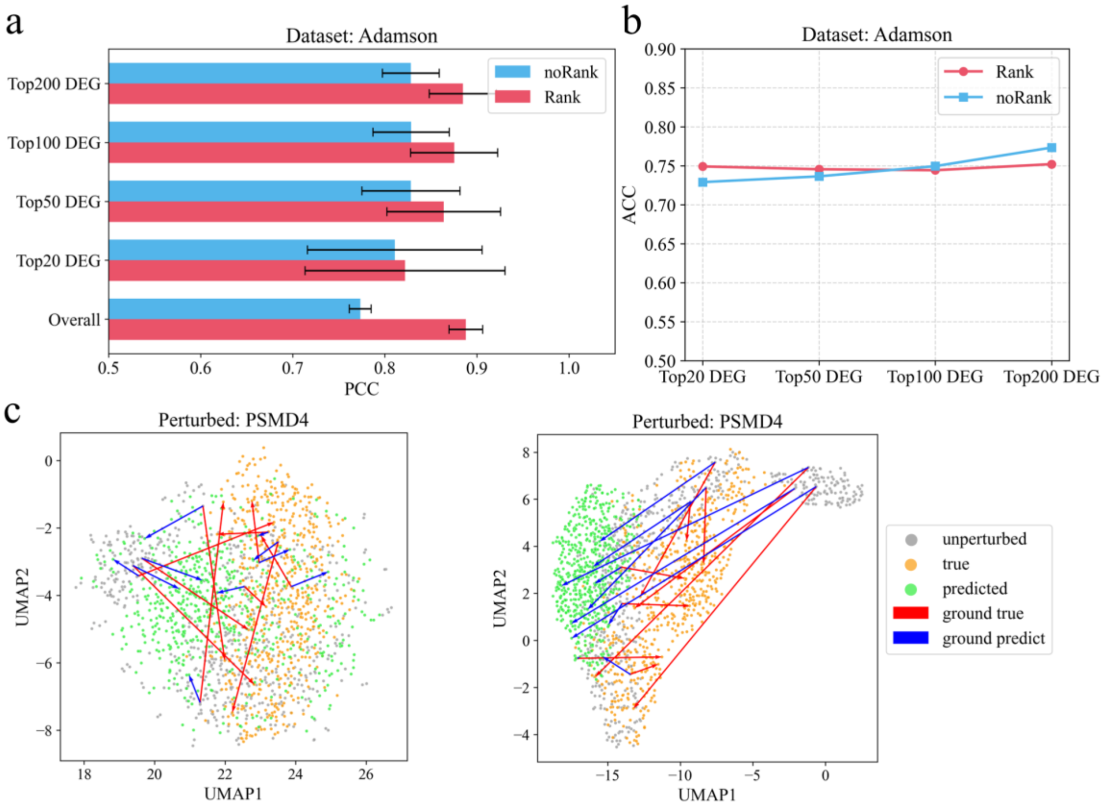


**Supplementary Figure 2: Evaluation of scRGP module performance without the Rank module. a.** Pearson correlation coefficient of the model on the Adamson dataset. **b.** Accuracy of predicting the direction of gene expression changes on the Adamson dataset. a-b. 'Rank' denotes the scRGP model, and 'noRank' denotes the scRGP model without the Rank module. **c.** UMAP plots of unperturbed cells, true perturbed cells, and predicted perturbed cells following perturbation of PSMD4. Arrowheads originate from unperturbed cells: blue arrowheads point to predicted post-perturbed cells, and red arrowheads point to true post-perturbed cells. The left panel shows predictions from the model without the Rank module, and the right panel shows predictions from the scRGP model.
